# Choosing between Glycolysis and Cholera: identifying glycolipid- and enzyme-based markers of extracellular vesicles in non-model organisms

**DOI:** 10.64898/2026.11.16.767010

**Authors:** Mads Zippor, Jonas Klejs Hemmingsen, Anders T Boysen, Ilaria Bellini, Hannah Weissinger, Litten Sørensen Rossen, Francesca Loria, Natasa Zarovni, Serena Cavallero, Stefano D’Amelio, Bianka Raupach, Andrew R Williams, Jørgen Kjems, Kenneth A. Howard, Peter Nejsum, Bradley Whitehead

## Abstract

The lack of universal markers for extracellular vesicles (EVs) is delaying progress in the field, especially research using non-model organisms. To develop effective options for the detection of EVs, we explored glycolipid-based and protein-based markers.

In this study, we show that cholera toxin subunit B (CTxb), which targets the glycosphingolipid GM1 in vertebrates, also binds to EVs from various non-model organisms enabling detection. However, whilst CTxb selectively detect EVs from cell culture and parasite media, it is unsuitable for complex biofluids, such as serum and plasma due to cross-reactivity with high- and low-density lipoproteins.

Additionally, we show that the universally conserved glycolytic enzyme enolase is found in EVs from phylogenetically distant species. Multiple highly conserved regions exist across human and several representative non-model organism orthologues, enabling detection of EVs from non-model organism using antibodies against human enolase. Furthermore, the conserved enzymatic activity of enolase enables detection of EVs using an enzymatic assay.

Together CTxb and enolase provide a tandem approach for detection of EVs from non-model organisms by targeting different, conserved components of EVs.

## Introduction

The study of extracellular vesicles (EVs) is central for understanding intercellular communication as well as host–pathogen interactions[1, 2]. However, progress in the field is hindered by the heterogeneity of EVs, with the lack of cross-taxon universal markers being a bottleneck especially in non-model organism EV research[3, 4].

Detection of EVs commonly relies on well-characterized protein markers such as the tetraspanins CD9, CD63, and CD81[5]. Unfortunately, these are largely limited to mammalian systems[6] and species-specific. Furthermore, commercial antibodies targeting known EV markers in non-model organisms are lacking, meaning that *de novo* antibody generation would be required, which is both time-consuming and expensive. This hinders adherence to MISEV guidelines, specifically concerning the accurate quantification and characterization of EV populations[7] with one of the major limitations in non-model organism EV-research, such as research in helminth EVs, being the lack of markers that can distinguish between EVs and non-vesicular components[8].

Although several candidates have been investigated as universal markers, none have proven definitively successful. A prominent example is phosphatidylserine (PS), which is displayed on EVs but is also highly expressed on apoptotic cells and platelets[9]. However, its detectable presence across different EV populations is debated and dependent on the reagents used[10–13]. Furthermore, the abundance of PS is seemingly context-dependent, as seen with levels of PS^+^-EVs being used as biomarkers for various cancers[14–16] as well as being associated with aortic stenosis, systemic lupus erythematosus, and thrombotic events[17, 18]. Crucially, in the context of non-model organisms, further limitations arise. For instance, some parasitic helminths express PS on the tegument, and tegument fragments are a potential contaminant in helminth EV preparations[19]. In contrast, PS is not widely found in bacterial outer membrane vesicles (OMVs)[20–22], highlighting its unsuitability as a universal EV marker.

With the goal of finding new candidates for universal markers enabling cost-effective detection, quantification and characterization of EVs and thereby addressing the challenges of non-model organism EV-research, we focused on two distinct classes of conserved molecules: the glycosphingolipid monosialotetrahexosylganglioside (GM1) and the glycolytic enzyme enolase.

GM1 is highly enriched in the membrane of human EVs compared to whole-cell membranes, which can be exploited for selective targeting of EVs[23, 24]. This enrichment might be attributed to vesicle biogenesis, as GM1 is abundant in lipid rafts[25], which are implicated in the formation of EVs and persist as well documented components of the resulting vesicles[26, 27]. Of note, lipid rafts have long been targets for detecting EVs, albeit indirectly, for example the commonly used EV marker flotillin-1, being a component of lipid rafts[5, 28].

In humans, GM1 on EVs has been targeted using cholera toxin subunit b (CTxb)[29, 30]. This interaction reflects the function of CTxb in *Vibrio cholerae* infection, where it mediates the cellular uptake of cholera toxin via a highly selective binding to GM1[31]. However, CTxb has not been used to target EVs from non-model organisms, where the presence of GM1 remains poorly characterized.

While GM1 serves as a lipid-based marker, enolase offers a protein-based alternative. Enolase fits the criteria of a highly conserved protein[32, 33], which would be expected for it to found in EVs from phylogenetically distant species. Such highly conserved proteins are often involved in fundamental cellular processes, such as DNA packaging[34] or glycolysis[35]. However, unlike prevalent mammalian EV proteins such as tetraspanins, TSG101, and ALIX, which play established roles in EV biogenesis and cargo loading [36, 37], a specific functional role has not been established for enolase despite its ubiquity.

To evaluate CTxb targeting GM1 and enolase-based methods as universal tools for detection of EVs, we assessed their presence across a broad biological spectrum and their specificity in distinguishing EVs from non-vesicular fractions and lipid contaminants.

Initially, to robustly validate the targeting of mammalian EVs with CTxb and its ability to distinguish EVs from non-EVs, human EVs genetically engineered with a fluorescent tag were used. This, importantly, enabled comparison of the labelling efficiency of CTxb with the known mammalian marker CD63 and the fluorescent EVs ensured a well-defined EV-population for comparison. Then to start evaluating the binding efficacy of CTxb across diverging species, we focused on the field of parasitic helminths, where EVs are studied extensively, as helminth EVs are presumed to mediate immune modulation of the host[2, 38–40]. However, tools for identification and characterisation of isolated EVs are limited and to date rely upon proteomics[8]. EVs from porcine roundworms *Ascaris suum* and *Trichuris suis* and the roundworm of marine mammals *Anisakis* spp. were investigated, and then to further extend the evaluation of cross-domain applicability, OMVs was also included.

Regarding enolase, a strong sequence homology across increasingly diverging species as well as a presence in EVs from these species were confirmed. Antibodies against human enolase with immunogens within highly conserved regions were tested for detection of EVs from *A. suum* and *T. suis*. Furthermore, to complement this antibody-based approach, an orthogonal method, utilizing the enzymatic activity of the enolase and therefore species agnostic, was evaluated.

Hereby, we aim to provide new methods for EV-detection in non-model organism taking advantage of glycolipid- and protein-based detection strategies, specifically, with the goal of enabling detection of EV-containing fractions after isolation, thereby confirming the successful enrichment of vesicles.

## Materials and Methods

### Engineering of mCherry-EV generating cells

An mCherry fused to a *686-truncated PTGFRN scaffold with a 3x SGGGG linker was inserted into the genome of HEK293T using transposon technology, as previously described[41]. Briefly, on the day before transfection, 300,000 HEK293T cells were seeded in a 6-well plate. 4 μg of donor plasmid and 1 μg of hyperactive piggyBac plasmid DNA were complexed with 15 μg of PEI Max, which was incubated for 30 min at room temperature and then added dropwise to the cells. The cells were incubated overnight at 37°C and 5% CO_2_. The following day, the media were exchanged, and engineered cells were selected using 150 μg/mL hygromycin B. The cells were maintained in selection media until a pure engineered population was achieved.

### Cell culture and EV-isolation

RAW 264.7 (TIB-71-ATCC) and HEK293 cells were cultured in DMEM plus 10% fetal bovine serum (FBS) with penicillin and streptomycin (P/S) at 37°C and 5% CO_2_. EV-depleted FBS was generated by diluting FBS 1:4 in DMEM and subjected to ultracentrifugation at 200,000 x g overnight, as previously described[42]. At 70% confluence RAW 264.7 media was washed twice in PBS prior to incubation in DMEM+5% EV-depleted FBS+P/S, whereas HEK293 cells were incubated in Optimem (ThermoFisher). Cell conditioned media (CCM) were harvested after 48 hours and sequentially cleared at 300 x g for 10 mins, 2000 x g for 15 mins and 10,000 x g for 30 mins prior to concentration using Pierce Protein Concentrators PES, 100K MWCO (ThermoFisher), followed by size exclusion chromatography (SEC).

### EV production from the nematodes *Ascaris suum, Trichuris suis* and *Anisakis* spp

Adult *A. suum* were sourced from naturally infected pigs at a commercial abattoir prior to washing and culture as previously described[43]. Adult *T. suis* were collected from experimentally infected pigs as previously described[44, 45] (Animal experimentation was approved by the Danish Experimental Animal Inspectorate (License number 2021-15-0201-00831)). *Anisakis* spp. (infective third stage larvae, L3) were isolated from the body cavity of *Clupea harengus* (FAO zone 27) and *Merluccius merluccius* (FAO zone 37) as previously described[46].

Briefly, nematodes were extensively washed in antibiotic and antimycotic solutions to prevent contamination, which was validated by endotoxin assays and agar plate growth assays as previously described[47]. After washing, nematodes were cultured for 3 days in RPMI in the presence of streptomycin (100µg/ml), penicillin (100U/ml), ciprofloxacin (5µg/ml) and amphotericin B (630ng/ml) at 37°C and 5% CO_2_. Conditioned media were harvested daily followed by clearance at 300 x g for 10 min, 2000 x g for 10 min prior to concentration using a centrifugal filter with a 10 kDa molecular weight cut-off (Merck), followed by SEC.

### Collection of outer membrane vesicles

*E. coli* DH5α with ampicillin resistance were used for OMV production. A bacterial stab was inoculated into LB supplemented with ampicillin and cultured overnight at 37°C with continuous shaking, in a total volume of 500 mL. Bacterial cells were removed by centrifugation at 3,000 × g for 20 minutes, followed by filtration (0.2 µm) using a Stericup vacuum filter. The supernatant containing OMVs was concentrated using Amicon Ultra centrifugal filters with a 100 kDa molecular weight cut-off. The resulting concentrate was then subjected to ultracentrifugation at 100,000 × g for 90 minutes at 4°C prior to washing with PBS and re-pelleting at 100,000 x g for 90 minutes.

### Plasma collection

Whole blood was collected into heparinised vacutainers (BD Bioscience) prior to immediate generation of platelet poor plasma via two sequential centrifugation steps at 3,200 x g for 15 minutes performed at room temperature to generate platelet poor plasma whilst minimising platelet activation. Plasma (500 µl) was then subjected to SEC and fractions were collected for analysis. Blood was obtained from 3 healthy anonymous volunteers attending the Danish blood bank at Aarhus University Hospital.

### Size exclusion chromatography

EVs were isolated by SEC using qEV 70 nm columns (iZON Science Ltd.), following the manufacturer’s protocol. 500 µL of sample was applied to the SEC column and eluted by adding sterile-filtered PBS. The EVs were collected in either fractions 7-9 or 7-10, as specified in the results, and kept at −80°C until use. Fractions between 10 and 24 were pooled as protein fractions, with the exact fractions used for each experiment stated in the results section.

### Nanoparticle tracking analysis

For both EV and protein enriched samples used for measuring enolase activity, the nanoparticle concentration and size distribution were determined by Nanoparticle Tracking Analysis (NTA) using Particle Metrix ZetaView PMX-120 (software version 8.05.12 SP) For each sample, two/three measurements were conducted in scattering mode, as previously described by Mladenović *et al.* [48].

For all other samples, than the ones used for the enolase activity assay, NTA measurements were performed using a NanoSight NS300, (Malvern Panalytical) with three 60-second videos captured and data analyzed using NTA 3.2.16 software.

### Cryo-transmission electron microscopy (TEM)

The samples used for cryo-TEM had a maximum of one freeze-thaw cycle and were thawed just prior to sample preparation. 3 µL of enriched EV-sample was dispersed to a glow-discharged, lacey carbon-coated 200 mesh copper grid (Quantifoil). Grids were then back-blotted with 1 µL of Milli-Q water added to the non-carbon film side, followed by blotting for 3 s on filter paper. The grids were then vitrified by plunging into liquid ethane (approximately −184°C) and subsequently stored in liquid nitrogen before imaging.

A Titan Krios G3i (Thermo Fisher Scientific) with a K3/BioQuantum detector/energy filter setup (Gatan) was used for data collection. The acceleration voltage was 300 kV. Scale bars were applied to images using ImageJ.

### Immuno-/ dotblots

For immuno- and dotblotting SEC-fractions or purified human LDL (ThermoFisher) or HDL (Sigma Aldrich) were immobilised on 0.45μm nitrocellulose membranes (BioRad) by spotting 3 µl and allowing to dry, prior to blocking with 5% (w/v) bovine serum albumin (BSA) in PBS for 30 minutes. Blots were then incubated with the indicated primary antibodies or biotinylated-CTxb (CTox-C34779, ThermoFisher) for one hour in 1% BSA PBS before washing in PBS + 0.05% Tween-20 for protein targeting antibodies or PBS for CTxb labelled blots. HRP conjugated secondary antibodies or streptavidin-HRP (RCD Systems) were then added for one hour followed by washing and development with Clarity Western ECL substrate using a Chemidoc Imager (Biorad). Primary antibodies used are as follows: anti-CD63 (10628D, ThermoFisher), anti-ApoB (MAB4124, RCD Systems), anti-ApoA1 (MAB36641, RCD Systems), anti-mCherry (ZRB2497, Sigma-Aldrich), anti-enolase (PA5-102334, ThermoFisher). Secondary antibodies used were HRP-conjugated anti-mouse secondary antibody (31432, Thermo Fisher) or HRP-conjugated anti-rabbit secondary antibody (656120, Thermo Fisher).

For HDL and LDL capture from human plasma anti-ApoA1 and anti-ApoB antibodies were spotted onto nitrocellulose membranes, allowed to dry prior to blocking with 5% (w/v) BSA in PBS and incubation with plasma diluted in 1% BSA PBS overnight at 4°C. Blots were washed and then incubated with biotinylated-CTxb prior to detection using streptavidin-HRP and development with Clarity Western ECL substrate using a Chemidoc Imager. Fucose inhibition assays were performed by spotting *A. suum* EVs or protein fractions on nitrocellulose membranes prior to blocking and incubation with biotinylated CTxb or UEA-I lectin (VectorLabs) in the presence or absence of 50mg/mL L-fucose followed by detection using Strep-HRP as per dotblots above.

Dot blot intensity was quantified using ImageJ v1.51 by measuring the integrated density of each dot, following background correction with a rolling ball radius set to 25 pixels.

### NanoFlow Cytometric analysis

Nanoflow cytometry analysis of EVs were performed using CytoFLEX nano (Beckman Coulter, Souzhou, China). QC were performed using CytoFLEX nano QC Scatterspheres (Beckman Coulter, C93495-AA) and CytoFLEX nano QC Fluorospheres (Beckman Coulter, C93496-AA). For HEK293T EV samples, data from 20,000 detected particles was recorded at a controlled flow of 1 μL/min, whereas for the *Anisakis* spp. EV samples the recording was done for 2 minutes at a controlled flow of 3 uL/min.

Anti-CD63 (MA5-18149, Invitrogen) and Cholera toxin subunit b-AF488 (2873181, Invitrogen) were used to label the HEK293T EV samples and mCherry-labeled HEK293T EV samples, while Cholera toxin subunit b-AF647 (C34778, ThermoFisher) was used for the *Anisakis* spp. EV samples. For both HEK293T EVs, mCherry-labeled HEK293T EVs, and *Anisakis* spp. EVs, 1 µL of the isolated EVs was incubated with 1 µL of one of the antibodies, resulting in a 1:1 dilution of the EVs. The mixture was then incubated for 30 minutes at 37°C. Next, for the HEK293T EV samples and the mCherry-labeled HEK293T EV samples 1 μL of the EV:antibody mix was diluted 1:1000 in PBS, resulting in the EV samples being diluted 1:2000. For the *Anisakis* spp. EV samples 1 μL of the EV:antibody mix was diluted 1:4000 in PBS, resulting in the EV samples being diluted 1:8000. The following control samples were included; unstained samples, PBS-only, and PBS + antibody, all of these were acquired at the same dilution as the stained samples.

The data was analyzed using FlowJo^TM^ v. 10.10 Software (BD Life Sciences). Based on a PBS-only control, background events from the buffer solution were excluded using an “inclusion” gate (VSSC1-H histogram). Then doublets were excluded using VSSC1-Width vs VSSC1-H. For the mCherry-labeled HEK293T EV samples, to ensure that only EVs positive for mCherry were included in the subsequent analysis, gating was based on regular HEK EVs and either the PBS + anti-CD63 or the PBS + CTxb, with no more than 1.0 % of events from these controls falling within the positive gate. The positive gates for CTxb and CD63 were then based on unstained mCherry-EVs with the same threshold of 1.0% (Supplementary Fig. 1). For the *Anisakis* spp. EV samples the gating was based on the unstained sample and the PBS + CTxb (Supplementary Fig. 2)

### Total internal reflection fluorescence microscopy

Total internal reflection fluorescence (TIRF) microscopy was performed using an Oxford Nanoimager (ONI) microscope. EVs were incubated with CTxb conjugated to Alexa Fluor™ 488 (Invitrogen, #2873181) for 30 min at 37°C. Subsequently, EVs were membrane-labeled and captured on a passivated surface using an established method[49]. In brief, EVs were mixed with 300 DOPE-ATTO (ATTO-TEC) and 150 biotinylated lipid-anchors (DSPE-biotin, ATTO-TEC)) per EV in PBS and incubated for 30 min at 37°C under gentle shaking. The biotinylated and fluorescently labeled EVs were then captured on a passivated glass slides with a microscope chamber (sticky-Slide VI 0.4, ibidi). The glass surface passivation was prepared as described previously[49]. In brief, they were cleaned, passivated, and coated with BSA-Biotin and subsequently covered with neutravidin. EVs were added to the surface and allowed to bind for 5 min, before being washed away using 2 mL PBS. The data was acquired with a 100×1.4 NA oil immersion objective and an exposure time of 100 ms. 162 images were acquired for each condition. Images in two channels were taken successively for each field of view, with the 488 laser at 7% and the 640 laser at 15%. One field of view corresponds to a physical field of view of 50 μm x 80 μm.

TIRF microscopy data was analyzed using an in-house developed automated python-based image analysis. Spots corresponding to EVs were detected as described before[49]. The membrane channel was used for spot detection and the background-corrected signal for the respective spots was extracted from both channels (membrane and CTxb). For the negative control, EVs without CTxb staining were detected and the corresponding CTxb signal was extracted from the control sample containing antibody in PBS without EVs.

### Enolase activity assay

The enolase activity of EV fractions 7-9 and protein fractions 16-24 isolated by SEC from *A. suum* were assessed by a fluorometric EV-Enolase kit (HansaBioMed Life Sciences; #HBM-K691-EN), following the manufacturer protocol. Standards, samples, blanks, and background controls were performed in duplicate on black non-binding polystyrene microplates (HansaBioMed Life Sciences). Fluorescence intensities were read in kinetic mode, using Tecan GENios Pro microplate reader (λex = 540 nm; λem = 590 nm).

### Multiple sequence alignment and 3D-mapping

To determine possible regions able to be targeted by commercially available antibodies, a multiple sequence analysis (MSA) was performed using EMBL Clustal Omega[50]. The results were visualized using Jalview 2.11.1.4[51].

A reference 3D structure of human alfa-enolase (P06733) was obtained from the Protein Data Bank. Using the ConSurf web server[52], the MSA-file obtained through EMBL Clustal Omega was utilized in determining and mapping conservation scores onto the 3D structure of the of human alfa-enolase. The results were visualized using PyMOL (Schrödinger, v3.0).

### Statistical analysis

Statistical comparisons were conducted in GraphPad Prism v10. Comparisons between two groups were carried out using either an unpaired, two-tailed t-test or a ratio paired t-test, whereas comparisons involving more than two groups were performed using one-way ANOVA, followed by Tukey’s multiple comparisons test, unless otherwise specified. A p-value < 0.05 was considered statistically significant.

## Results

### GM1 on EVs can be targeted by CTxb

EVs were isolated from cell conditioned media of genetically engineered human HEK293T-cells expressing mCherry via a PTGFRN scaffold on EVs (mCherry-HEK-cells) and of murine RAW264.7 cells using SEC to investigate the binding of CTxb to EVs.

For the mCherry-HEK-cells, SEC fractions 7-9 (F7-9) and 10-15 (F10-15) were collected as EV-enriched fractions and protein-enriched fractions, respectively. Cryo-TEM was used to confirm enrichment of vesicular structures (Fig. 1A), and vesicle size distributions were determined using NTA (Fig. 1B). Dotblots of the fractions showed substantial CTxb-, CD63-, and mCherry-signals in F7-9 with minimal signal in F10-15, except for mCherry (Fig. 1C-D). The mCherry signal in F10-15 was likely due to free mCherry in the cell culture media being eluted in the protein fractions, which the measurement of fluorescence intensity of the all the eluted fractions clearly showed (Fig. 1E). Taken together, this indicated the binding of CTxb to mCherry-HEK-EVs.

**Figure 1.**
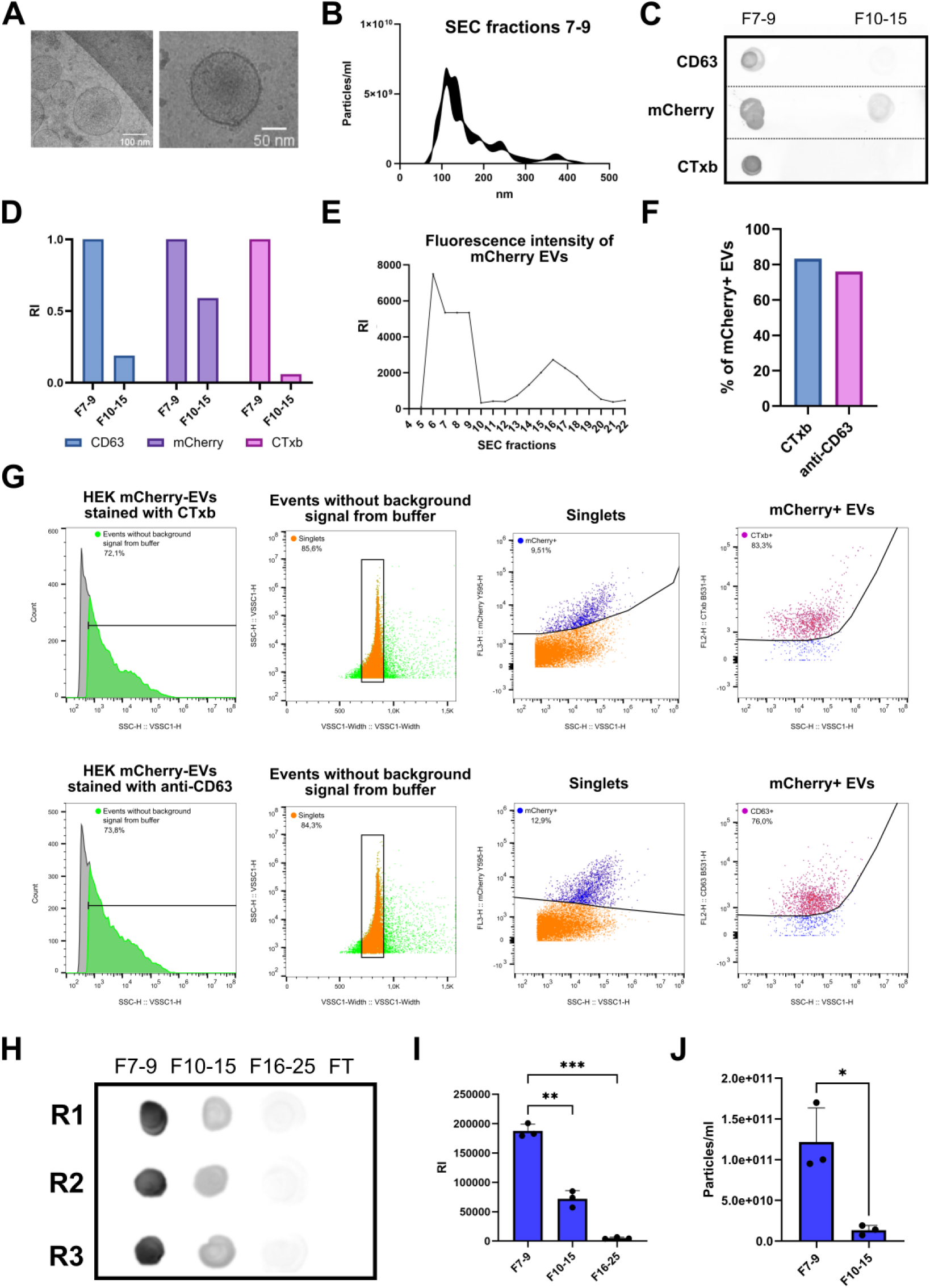
**A)** Cryo-TEM images of HEK-mCherry-EVs (scalebar, left:100 nm, right: 50 nm). **B)** Particle size distribution of F7-9 isolated by SEC from HEK mCherry-cells estimated by NTA. **C)** dotblot of F7-9 (EVs) and F10-15 (proteins) isolated by SEC from HEK mCherry-cells using CTxb, anti-mCherry, and anti-CD63. **D)** Densitometric quantification of the dotblot in C with intensities normalised to highest value for each blot. **E)** Fluorescence intensity of the SEC-fractions isolated from HEK mCherry-cells. **F)** Percentage of mCherry+ EVs positive for CTxb or anti-CD63 when analysed by nano-flow cytometry **G)** Gating performed for nano-flow cytometric analysis of HEK-mCherry-EVs stained with either CTxb or anti-CD63. **H)** dotblot of F7-9 (EVs), protein fractions F10-15 and F16-25, and flow through (FT) of the SEC isolated RAW264.7-EVs using CTxb. Performed in three replicates. **I)** Densitometric quantification of the dotblot in G. **J)** NTA-based particle quantification of the F7-9 and F10-15 isolated by SEC from RAW264.7 cells. RI; relative intensity.

CTxb binding to mCherry-HEK-EVs was further confirmed using nano-flow cytometry (Fig 1G). The F7-9 were stained with either CTxb conjugated to Alexa Fluor™ 488 or Alexa Fluor™ 488-conjugated anti-CD63. On average 11.2% of the particles in F7-9 were positive for mCherry, suggesting around one in ten particles detected by nano-flow cytometry were EVs. Both GM1 and CD63 were successfully detected on the EVs with 83.3% of the mCherry-positive EVs being labelled with CTxb and 76.0% being labelled with anti-CD63 (Fig 1F). This supported the evidence from the dotblots and confirmed the binding of CTxb to individual EVs. Furthermore, the percentage of EVs labelled with CTxb was similar to the percentage labelled with anti-CD63, demonstrating GM1 as a viable marker for human EVs.

CTxb was also shown to label EVs derived from murine RAW264.7 cells. F7-9 (EVs), F10-15 (proteins), F16-25 (proteins), and flow through were collected and dotblotted with CTxb, showing the most intense staining in F7-9 followed by minimal staining in F10-15 (Fig 1H-I).

### CTxb labelling of non-model organism derived EVs

EVs from the nematodes *A. suum, Anisakis* spp., and *T. suis* isolated using SEC and *E. coli* OMVs isolated by ultracentrifugation were used to investigate the CTxb-based detection of EVs from non-model organisms.

The presence of EVs was validated by Cryo-TEM and (Fig. 2A, - D) and NTA (Fig. 2E) and EVs from all four species were successfully labelled using biotin conjugated CTxb. For the three nematode species, dotblots showed a high signal in the EV fractions with minimal cross reactivity in the non-EV fractions (Fig. 2B-C). For OMVs the dotblot showed labelling only in the presence of CTxb (Fig. 2B+F). Notably, the binding of CTxb to *A. suum* EVs was reduced by excess L-fucose present in CTxb labelling solution (Supplementary Fig. 3).

**Figure 2.**
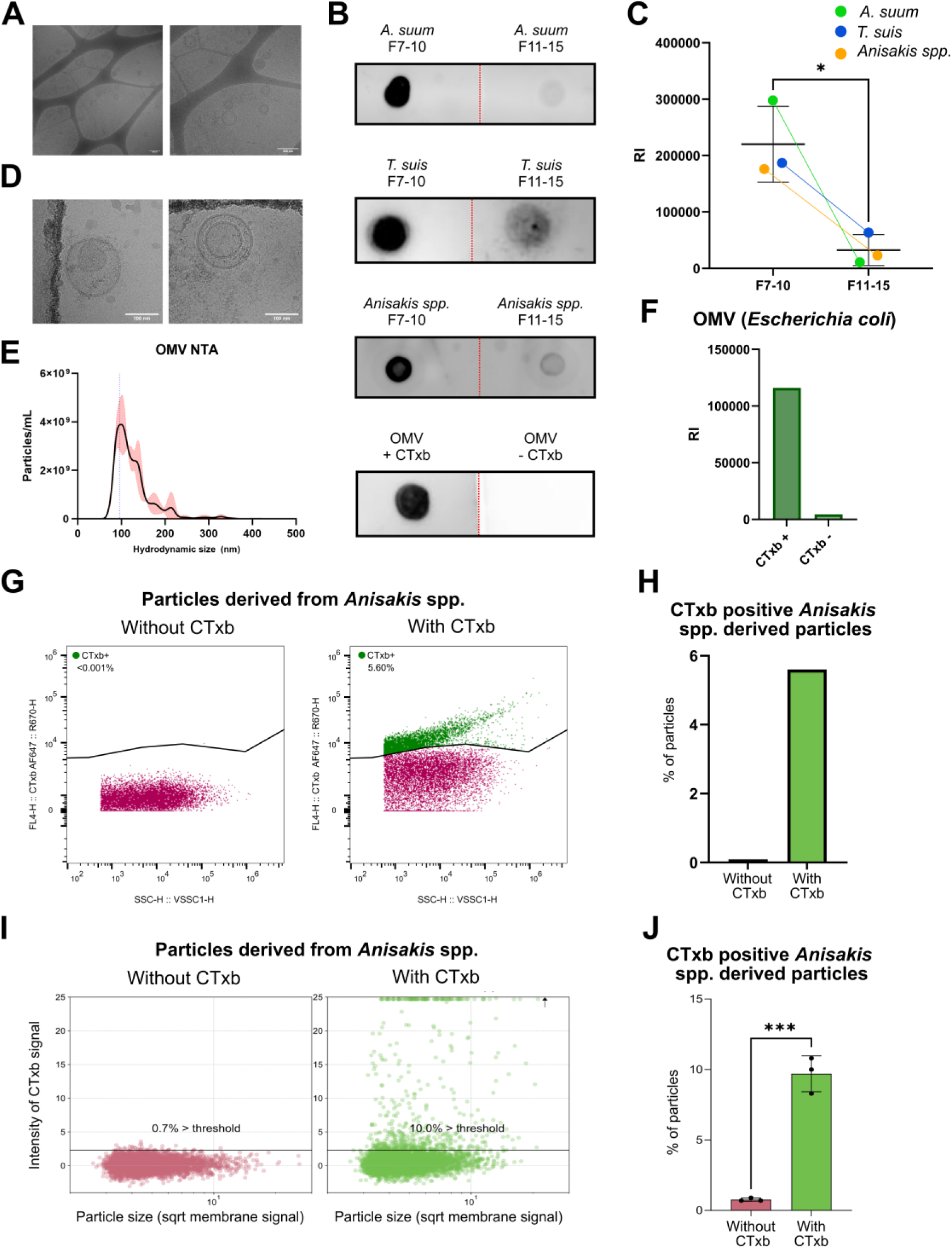
**A)** Cryo-TEM image of *Ascaris suum* EVs (scalebar, 200 nm). **B)** CTxb dotblots of EVs (F7-10) and proteins (F11-15) isolated from *A. suum*, *Anisakis* spp. or *Trichuris suis* by SEC. Also an dotblot of OMVs isolated by ultracentrifugation with and without CTxb. **C)** Densitometric quantification of the dotblots of *A. suum, Anisakis* spp., and *T. suis*. **D)** Cryo-TEM image of *Anisakis* spp. EVs (scalebar, 100 nm). **E)** Particle size distribution of OMVs estimated by NTA. **F)** Densitometric quantification of the OMV dotblot. **G)** Gating performed for nano-flow cytometric analysis of *Anisakis* derived particles (F7-10) stained with CTxb. **H)** Percentage of *Anisakis* derived particles positive for CTxb when analysed by nano-flow cytometry. **I)** TIRF microscopy analysis. Scatterplot showing the intensity of CTxb signal for individual *Anisakis* spp. derived particles. Threshold was set to 2.5 * standard deviation of the CTxb signal of particles without CTxb. Each condition contains 5290 spots. Spots outside of the axis range were visualized at the plot outline. Depicted scatterplot represents one out of three independent experiments. **J)** Quantification of CTxb-positive *Anisakis* spp. derived particles across three individual experiments. *** p-value < 0.001. RI; relative intensity.

To further examine the binding of CTxb to EVs from a non-model organism, TIRF microscopy was performed on immobilised, membrane labelled *Anisakis* derived particles (F7-10) stained with CTxb (Supplementary Fig. 4). Hereby, it was determined that 9.7±1.3% of the immobilised particles were CTxb-positive (Fig. 2I-J). The binding of CTxb to individual *Anisakis* derived particles was validated using nano-flow cytometry, which showed 5.6% of the total particles being CTxb-positive (Fig. 2G-H). The proportion of labelled particles was similar to the proportion of mCherry-positive particles in the HEK-mCherry-EV fractions identified using nano-flow cytometry (Fig. 1G), indicating that CTxb binds a large fraction of the *Anisakis* derived EVs.

These results highlighted the potential use of CTxb for the detection of EVs from non-model organisms.

### CTxb cannot be used to selectively detect plasma EVs

Next, we investigated if CTxb could be used to detect EVs in a complex biofluid such as plasma.

Dotblots for CTxb showed staining of SEC fractions typically enriched in EVs (Fig. 3A). To assess potential cross-reactivity to particles in plasma, we then tested CTxb labelling of low-density lipoprotein (LDL), and high-density lipoprotein (HDL), which are common contaminants of EVs preparations from plasma[53, 54] (Fig. 3B). Both pure LDL and pure HDL showed strong labelling with CTxb, suggesting a limitation of using CTxb to detect EVs isolated from plasma by SEC alone. To rule out potential co-isolation of EVs during LDL or HDL purification, samples were counterstained with anti-CD63, revealing no EVs in the LDL or HDL preparations, indicating binding of CTxb to the lipoproteins (Fig. 3B). As an orthogonal demonstration of CTxb binding of HDL and LDL we performed immunocapture of these lipoproteins using anti-ApoA (for HDL) and ApoB (for LDL) followed by detection of HDL/LDL using CTxb (Fig. 3C).

**Figure 3.**
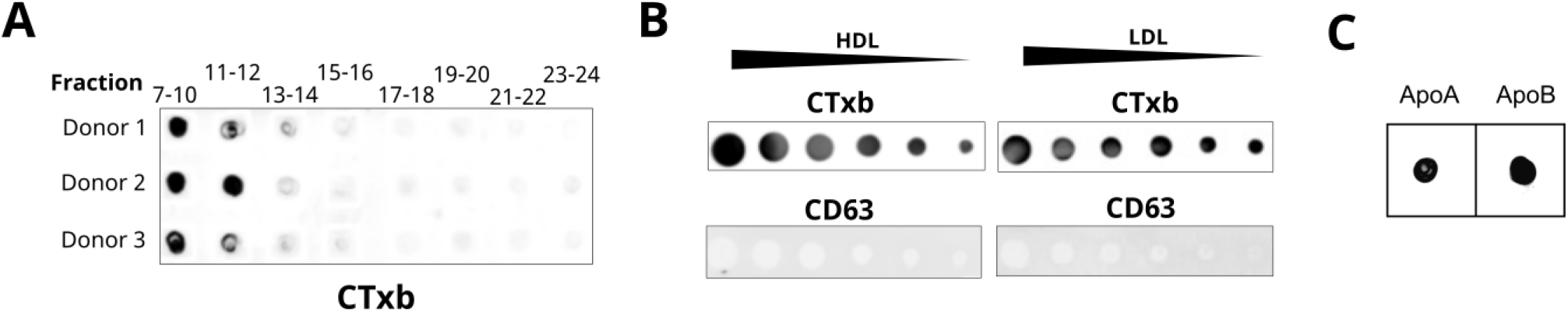
**A)** Human plasma (n=3) separated by SEC and dotblotted with CTxb. **B)** Purified HDL and LDL preparations dotblotted with CTxb and counterstained with anti-CD63 to confirm the absence of CD63-positive EVs. **C)** Immunocapture of ApoA and ApoB from human plasma prior to detection with CTxb to confirm selective reactivity with HDL and LDL, respectively.

These findings showed that CTxb is not suitable for EV detection or capture from complex biofluids, such as plasma or serum, unless complete removal of lipoproteins is confirmed.

### Enolase is a well-conserved EV-protein across multiple taxa

To expand the EV detection toolkit, we explored protein markers, which could provide methods for EV detection beyond a glycolipid-based method.

Using a 30 % lower boundary for sequence percentage identity, the EV-proteome of *A. suum* from Hansen *et al.*[6] was compared with the EV-proteomes from two other nematodes, namely *Trichuris muris* described by Eichenberger *et al.*[55] and *Brugia malayi* described by Harischandra *et al.*[56]. Only eight EV-proteins had orthologues present in EVs from the three analysed nematode species (Supplementary Fig. 5). The comparison was then expanded to the platyhelminth *Schistosoma mansoni* using the EV-proteome from Samoil *et al.*[57]. Of the eight proteins, only enolase was also identified in *S. mansoni* EVs. To further validate the presence of enolase in EVs from increasingly phylogenetically distant species, EV-proteomes from the protozoan parasite *Plasmodium falciparum*[58] and the prokaryote *Vibrio cholerae*[59], where CTxb is derived from, were investigated, and enolase was also found be present in EVs from these organisms.

Importantly, enolase appeared to be well conserved between the analysed species, with the lowest sequence identity between the species being 49.77% and the species most divergent from the others being *V. cholerae* (Fig. 4A).

**Figure 4.**
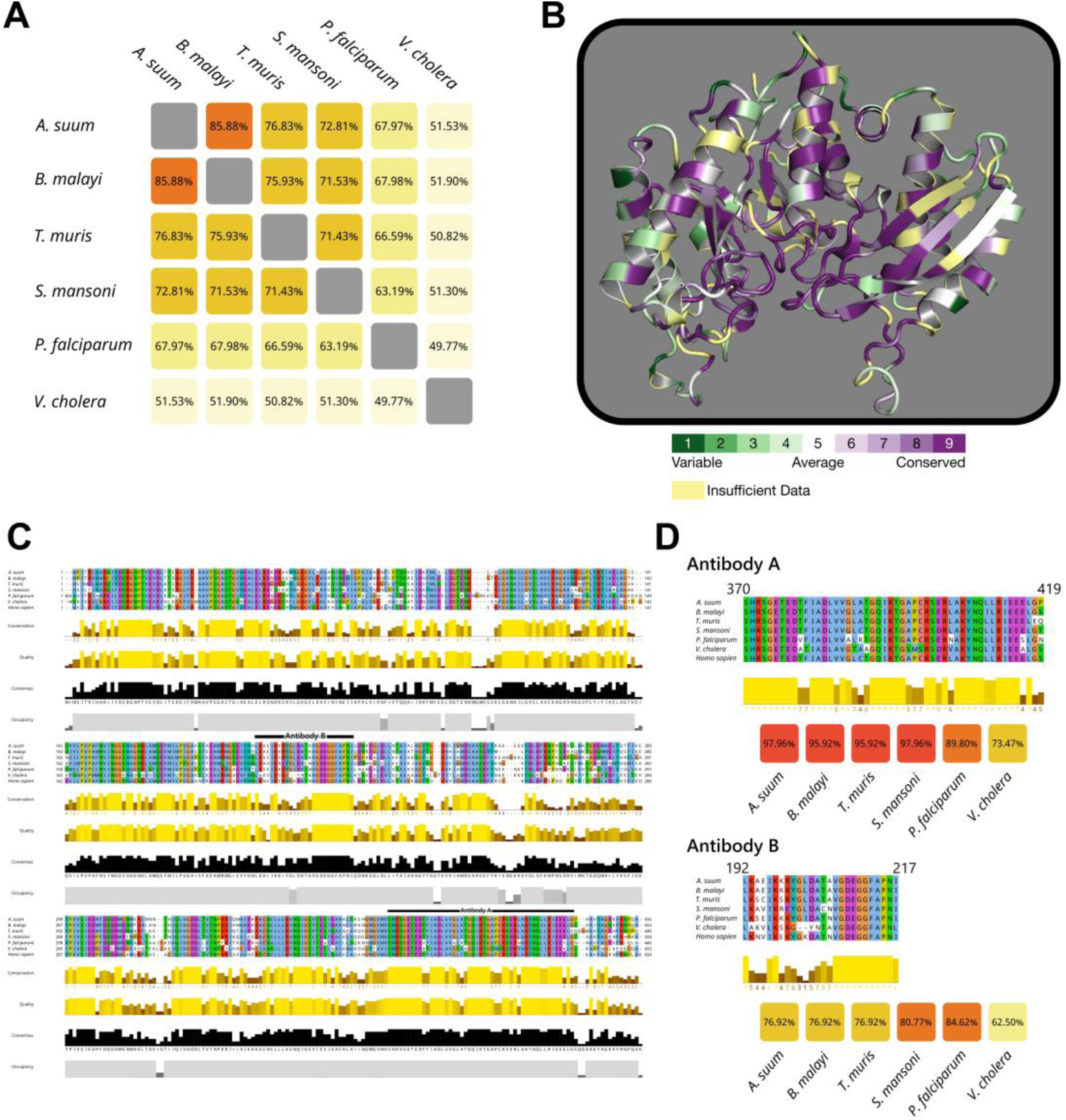
**A)** Percent identity matrix showing pairwise sequence similarity between enolase orthologues from *Ascaris suum* (UniProtID: E5D7U6), *Trichuris muris* (A0A5S6QD18) *Brugia malayi* (A0A0H5S082), *Schistosoma mansoni* (Q27877), *Plasmodium falciparum* (Q8IJN7), and *V. cholerae* (Q9KPC5). **B)** The 3D structure of the human alfa-enolase (P06733) with mapping of conservation scores based on the *A. suum*, *T. muris*, *B. malayi*, *S. mansoni*, *P. falciparum*, and *V. cholerae*. **C)** Multiple sequence alignment of the *A. suum*, *B. malayi*, *T. muris S. mansoni*, *P. falciparum*, *V. cholerae*, and human alfa-enolase with highly conserved regions across the species colored in the brightest yellow. **D)** Sequence of the immunogens for Antibody A and Antibody B aligned against the same sequence in enolase orthologues from *A. suum*, *B. malayi*, *T. muris S. mansoni*, *P. falciparum*. Sequence similarity between the sequences in these enolase orthologues and the immunogen is shown below each alignment.

Due to the lack of antibodies for non-model organisms available, we explored the potential use of commercially available antibodies against human alpha-enolase for detection of the investigated orthologues. Multiple sequence alignments were performed to evaluate if the orthologues had conserved regions suitable for binding of antibodies against the human orthologue. Several conserved regions of sufficient length to bind an antibody were identified (Fig. 4B-C). For two of these regions, we were able to find commercial antibodies raised against immunogens that are encompassed in the highly conserved sequences: Antibody A (PA5-102334, ThermoFisher) and Antibody B (HPA068721, Sigma) (Fig. 4D).

### The presence of enolase can be used to identify EVs from non-model organisms

Immunoblots of *A. suum* and *T. suis* were performed with the two commercially available antibodies against human alpha-enolase to investigate the binding to orthologues from non-model organisms. Antibody B had insufficient affinity for *A. suum* enolase and *T. suis* enolase to effectively distinguish EV fractions from non-EV fractions. Antibody A, however, was successful in distinguishing EV fractions (F7-10) from non-EV fractions (F11-15) (Fig. 5D-F).

**Figure 5.**
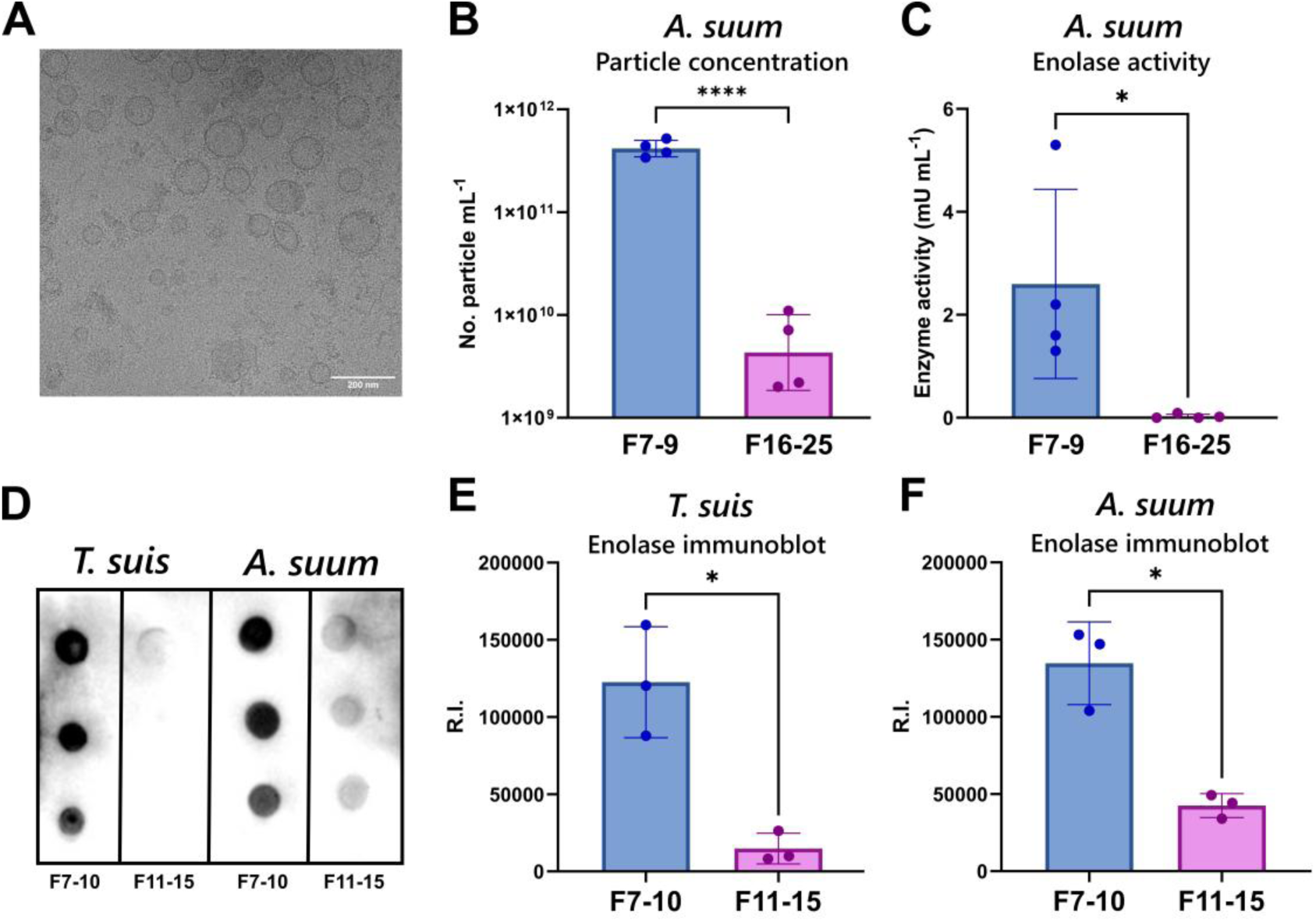
**A)** Cryo-TEM image of *A. suum* EVs (scalebar, 200 nm). **B)** Particle concentrations of F7-10 and F16-25 measured by NTA. **** p-value < 0.0001. **C)** Enolase enzyme activity of EVs (F7-10) and proteins (F16-25) isolated from *A. suum* by SEC. * p-value < 0.05. **D)** Immunoblot of F7-10 (EVs) and F11-15 (proteins) isolated by SEC from *A. suum* or *T. suis* using anti-enolase (Antibody A, PA5-102334 ThermoFisher). **E)** Densitometric quantification of the anti-enolase immunoblot of *T. suis*. **F)** Densitometric quantification of the anti-enolase immunoblot of *A. suum.* RI; relative intensity.

As an orthogonal method for detecting the enolase content, we measured the enzymatic activity of enolase. EVs (F7-9) and proteins (F16-25) isolated by SEC from *A. suum* were analysed using an enolase activity assay. In F7-9, the enolase activity was detectable at 2.4 mU/mL and showed significantly higher readouts than the activity in F16-25, which was barely measurable (Fig. 5C).

This highlights the potential of enolase as an EV-marker in non-model organism, with it being detectable either through cross-reactive antibodies targeting the human orthologue or via its intrinsic enzymatic activity.

## Discussion

In this study, we assessed the targeting of GM1, a glycosphingolipid consistently found in vertebrates[60], as well as an evolutionary conserved enzyme, enolase, for the detection of EVs from non-model organisms. Thereby, beginning to address the lack of reliable and broadly expressed EV-markers in taxa such as helminths[3, 8], bacteria[61], and plants[62].

We showed that CTxb binds to EVs and could be used for detecting EVs from different helminth species as well as OMVs. The proportion of CTxb-labelled particles ranged from 6% to 10% across nano-flow cytometry and TIRF microscopy. These values are comparable to the mCherry-positive percentage observed in HEK-mCherry-EV-fractions, which likely represent the actual proportion of EVs; notably, 83.3% of these were also CTxb-positive. Naturally, the EV proportions will depend on the starting material, isolation protocol, and the analytical method employed; here, TIRF specifically enriches for lipid particles, while nanoflow cytometry provides a non-selective analysis of the total particle population with the detection thresholds also differing between the two techniques. Nevertheless, the close alignment between CTxb labelling and mCherry expression would suggest that the majority of the vesicular population is effectively targeted by CTxb given the concentrations of “background” particles are similar.

Furthermore, the apparent surface binding of CTxb indicated its potential to be used for EV-capture across a broad range of species, possibly adding to a growing number of lipid-based separation methods[63]. However, detection of EVs was not possible in more complex biofluids such as serum and plasma, where CTxb also binds HDL and LDL, which could be due the presence of GM1 on these particles, with lipoproteins having been shown to bind and transport both exogenously and endogenously glycosphingolipids[64–66]. Therefore, without complete depletion of HDL and LDL we advise against the use of fluorescent CTxb for analysis of plasma EV preparations, e.g. nano-flow cytometric applications. Importantly, this finding does not interfere with identification of EVs from cell culture or parasite cultures, where CTxb presents a valuable tool for the discrimination of EVs from non-vesicular components.

The mechanism of CTxb binding to EVs from non-model organisms may differ from that in humans and mice. In humans, CTxb has been used for EV-labelling [29, 30], taking advantage of the clear enrichment of GM1 in human EVs compared to cell membranes[24]. While, GM1 and the enzymes needed for synthesis of gangliosides are highly conserved amongst vertebrates[60, 67], GM1 glycolipids have not been found in invertebrates[68], making it unlikely that GM1 mediates the binding of CTxb to EVs from the non-model organism investigated in this study.

It should be noted, however, that helminths with a blood stage in their life cycle have been shown to incorporate host blood group antigens [69, 70] and *Anisakis* spp. has been reported to incorporate fluorescent lipids into EVs *in vitro*[46], suggesting the possibility that glycosphingolipids such as GM1 may be acquired from the host environment. While this could be a potential mechanism for the helminths investigated, this is not the case for the *E. coli*-derived OMVs, as the *E. coli* from cultures performed in this work have never been exposed to a host environment.

Alternatively, the CTxb binding to EVs might be mediated by fucosylated oligosaccharides. CTxb has been shown to also bind L-fucose as well as fucosylated oligosaccharides, such as the histo-blood group antigen Lewis^x^[71, 72], providing a potential alternative binding site on EVs. Helminths, including the species investigated in this study, produce fucosylated glucans[73], with EVs from *S. mansoni* displaying fucose-containing glycans in the form of Lewis^x^[74, 75]. Indeed, we showed that the presence of L-fucose inhibits CTxb binding to *A. suum* EVs (Supplementary Fig. 3). Given that L-fucose and GM1 bind to different sites on CTxb[76], this finding indicates L-fucose as the specific target of CTxb binding on *A. suum* EVs, rather than GM1. Although not being investigated for all non-model organisms included in this study, the restriction of GM1 to vertebrates suggests the CTxb target in all these species is likely L-fucose or fucosylated oligosaccharides Further supporting this fucosylated glycans are also found across bacterial species[77] and LPS, which is present on the surface of OMVs, have in some bacteria been shown to contain L-fucose likely as a mean of immune evasion through molecular mimicry[78]. Adding to this, the carbohydrate mimicry between lipooligosaccharides of *Campylobacter jejuni* and human GM1 are important in the pathogenesis of Guillain–Barré syndrome and CTxb binds strongly to these lipooligosaccharides[79]. Likewise, CTxb-binding to LDL and HDL in our study could, besides the display of GM1, be due to fucosylated glycans on the surface of these particles[80], potentially pointing towards a broader limitation of CTxb in distinguishing EVs from other particles containing fucosylated glycans.

Irrespective of the CTxb binding mechanism to EVs from helminths, CTxb showed selectivity for EVs when assessed by dotblotting, nano-flow cytometry and TIRF microscopy. If binding proceeds via L-fucose we would predict significant binding in protein fractions from SEC, given the higher protein concentration in these fractions compared to the EV fractions. The pentavalent structure of CTxb may reconcile the specificity observed for EVs. This structure enhances avidity through multivalent interactions as is expected in an EV membrane or a lipid raft where target antigens are clustered and, therefore, may explain the increased binding of CTxb to EV presented L-fucose compared to that on secretory glycoproteins. Indeed, multivalent glycan-based inhibitors of CTxb increase effectivity by over 440-fold compared to monovalent forms[81, 82] and increased concentration of glycans in a model glycocalyx increased affinity of CTxb by approximately 11,000 fold[83]. Supporting this, we show that the fucose binding lectin UEA-I bound equally to *A. suum* EVs and protein fractions whereas this is not the case for CTxb despite both UAE-I and CTxb binding being inhibited by L-fucose, further implicating multivalent binding of CTxb to antigens in EV microdomains (Supplementary figure 6). The enhanced avidity conferred by multivalency may also explain the cross reactivity of CTxb to HDL and LDL, as both of these are fuscosylated[80], whilst not showing significant reactivity to free plasma proteins, where fuscosylated proteins are abundant. Taken together, this suggest that CTxb selectivity is driven by the spatial clustering of target glycans rather than their mere presence. Furthermore, this may present a novel angle to targeting EVs as nano-clustering of antigens on EVs may increase specificity of targeting ligands compared to free monovalent epitopes.

As a complementary approach to the glycolipid-based CTxb-detection of EVs, we demonstrated the detection of non-model organism derived EVs based on their enolase content both through cross-reactive antibodies targeting the human orthologue and via its intrinsic enzymatic activity. Enolase is well conserved and has been found in EVs from phylogenetically distant species[55–59], highlighting the possibility for a broader applicability of detection based on enzymatic activity. Sequence alignment of the immunogen of an anti-enolase from ThermoFisher (PA5-10233, Antibody A) showed a high degree of conservation across taxa, also demonstrating the potential of universal antibody-based detection upon selection of the correct immunogen. Still, this might not be feasible due to epitope divergence and the highly conserved enzymatic function of enolase might be the best option for detection of EV enriched fractions [33].

The reason for enolase’s conserved presence in EVs is not apparent. Multiple proteins were found to have orthologues in EVs from the nematodes investigated, however, of these, only enolase was identified in EVs from the platyhelminth *S. mansoni*. Furthermore, enolase was also found when the comparison was expanded to *P. falciparum,* and *V. cholerae*. The presence of enolase in EVs across distant species is consistent with enzymes involved in glycolysis usually being highly conserved[35]. However, other glycolytic enzymes, glyceraldehyde 3-phosphate dehydrogenase and malate dehydrogenase, present in EVs from all investigated nematodes, were not observed when the comparison was expanded to *S. mansoni*, *P. falciparum,* and *V. cholerae*. This raises the question why specifically enolase seems to be a conserved EV-protein.

While enolase is classically thought of as an enzyme involved in glycolysis, catalysing the conversion of 2-phosphoglycerate to phosphoenolpyruvic acid, additional functions of enolase have been uncovered. Enolase has been shown to have binding capacity for different nucleic acid types such as mRNA[84]. Furthermore, enolase functioning as a cell surface plasminogen receptor likely participating in pathogen adhesion to host cells, has been shown in bacteria such as *Streptococcus pneumoniae*[85], *Staphylococcus aureus*[86], and possibly also in the parasite *Giardia intestinalis*[87]. Intriguingly, EVs secreted by the liver fluke, *Fasciola hepatica* contain an enolase-orthologue, with anti-inflammatory properties and a potential role in mitigating liver fibrosis[88]. Given these diverse roles, we sought to determine whether enolase is intraluminal or associated with the surface in our EV populations. We conjugated Alexa Fluor™ 647 to the anti-enolase antibody for nano-flow cytometry analysis; however, this was unsuccessful. We cannot distinguish whether this is due to technical limitations of the antibody-fluorophore-conjugate or an intraluminal location of the protein. Thus, further work is necessary to establish the exact location of enolase in EVs.

The above-mentioned functions of enolase are cases of ‘moonlighting’. Proteins with moonlighting functions are multifunctional proteins, characterized by one polypeptide chain displaying distinct, physiologically relevant biochemical or biophysical functions without being a result of gene fusions or the presence of multiple proteolytic fragments[89]. These proteins are often highly conserved and typically expressed at high levels, with enzymes involved in sugar metabolism often presenting with numerous moonlighting functions[89, 90], which aligns with the observed multiple moonlighting functions of enolase.

## Conclusion

In this work, we show that CTxb and enolase can be used for the detection of EVs from several different non-model organisms. Thereby, providing a possible tandem method for the detection of EVs from non-model organisms by targeting different components of the EVs.

## Acknowledgements

This work was supported by the Danish Council for Independent Research (DFF-1032-00242B and 3105-00153B), by the Carlsberg Foundation (CF23-1020), and by the European Commission-funded H2020-FETOPEN-2016-2017 “The Extracellular Vesicle Foundry” (evFOUNDRY) project (Grant Agreement ID: 801367). The funders had no role in study design, data collection and analysis, decision to publish, or preparation of the manuscript. Flow cytometry was performed at the FACS Core Facility, Aarhus University, Denmark. The Cryo-transmission electron microscopy was performed at the National Cryo-EM Facility at Aarhus University, Denmark.

## Declaration of Interest Statement

Authors declare that they have no competing interests.

## Data Availability Statement

The data that support the findings of this study are available from the corresponding author upon reasonable request.

## Notes

### Competing Interest Statement

The authors have declared no competing interest.

