## Supplementary for "Choosing between Glycolysis and Cholera: identifying glycolipid- and enzyme-based markers of extracellular vesicles in non-model organisms"

#### Supplemental Figure 1

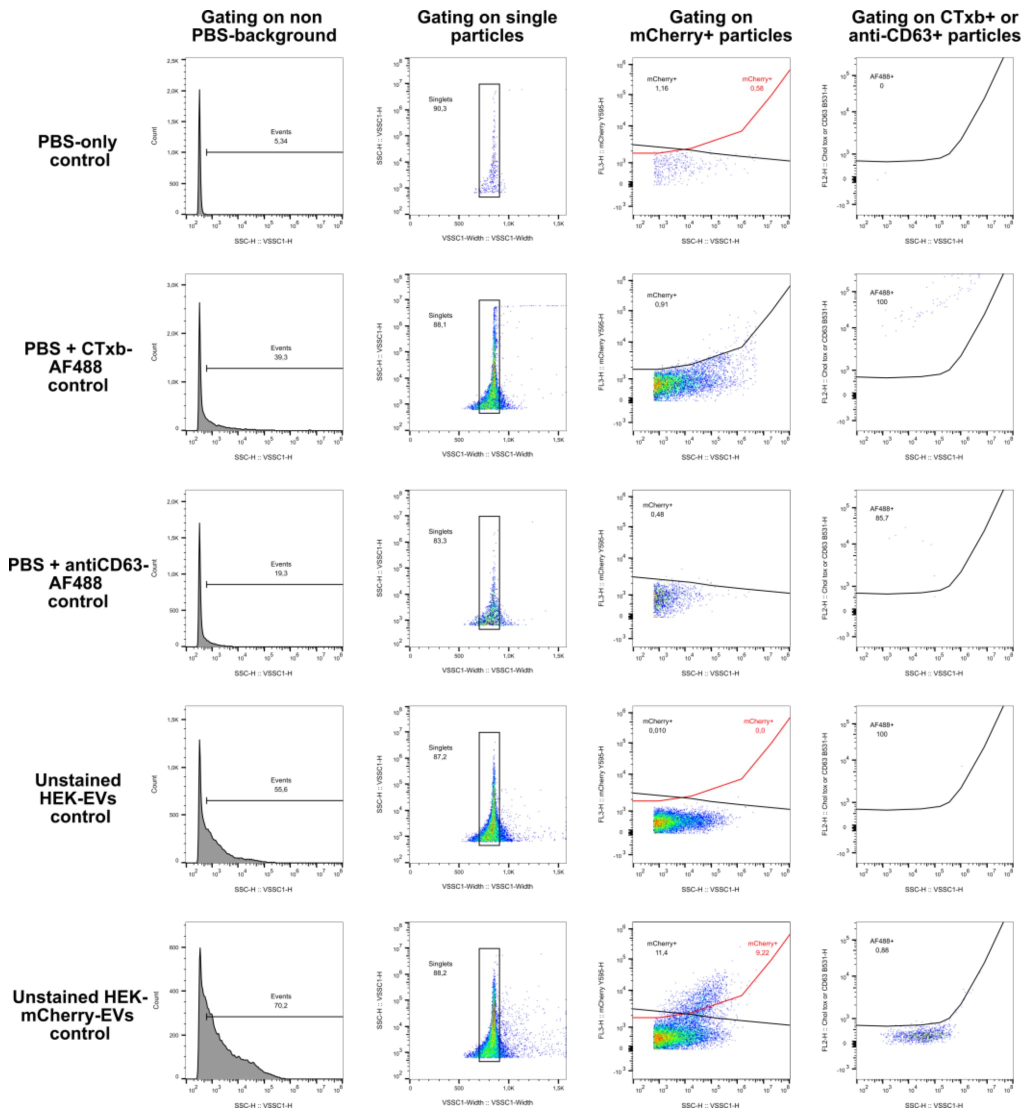


**Gating strategy for nano-flow cytometry on HEK-mCherry-EVs.** Based on a PBS-only control the background events from the PBS were gated out. Then based on Unstained HEK-EVs, PBS+CTxb-AF488 and PBS+antiCD63-AF488 the gating for mCherry-positive events were set with no more than 1.0 % of events from these controls falling in within the gate (On the figure the chosen gate for the CTxb stained sample is shown in red on the flow cytometry charts for PBS-only, Unstained HEK-EVs, and HEK-mCherry-EVs) The positive gates for CTxb-AF488 and anti-CD63-AF488 were then based on unstained HEK-mCherry-EVs with the same threshold of 1.0%.

**Supplemental Figure 2**

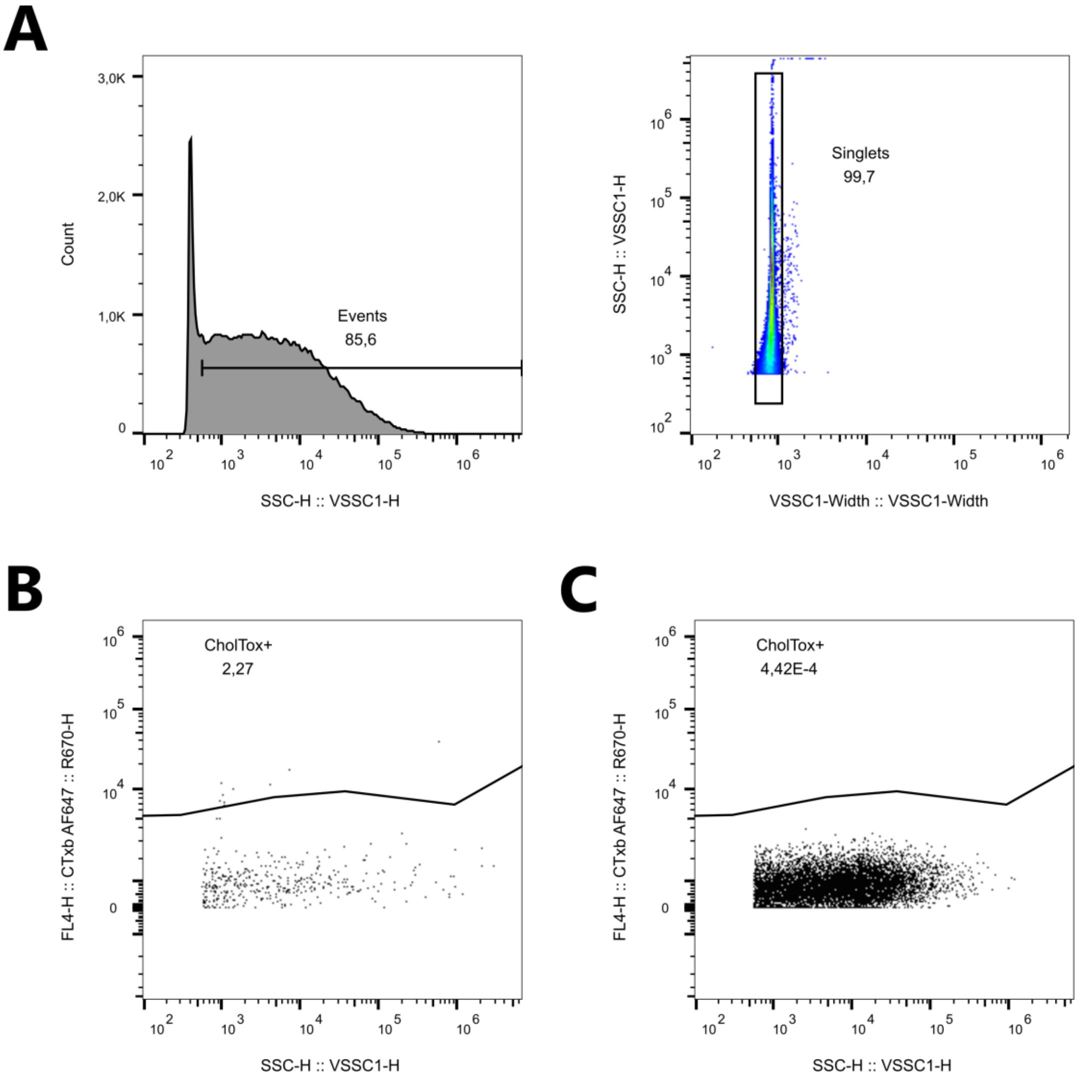


| | CTxb in PBS | Stained sample |
|---|---|---|
| Total events | 397 | 82 228 |
| CTxb-positive events | 9 | 4605 |

**Gating strategy for nano-flow cytometry on *Anasakis* derived EVs.** A) Based on a PBS-only control the background events from the PBS were gated out, followed by removal of dublets. The gates for CTxb-positve events where then set on: B) PBS + CTxb, which had only 397 events compared to the 82228 of the stained sample, and C) unstained *Anasakis* derived EVs.

**Supplemental Figure 3**

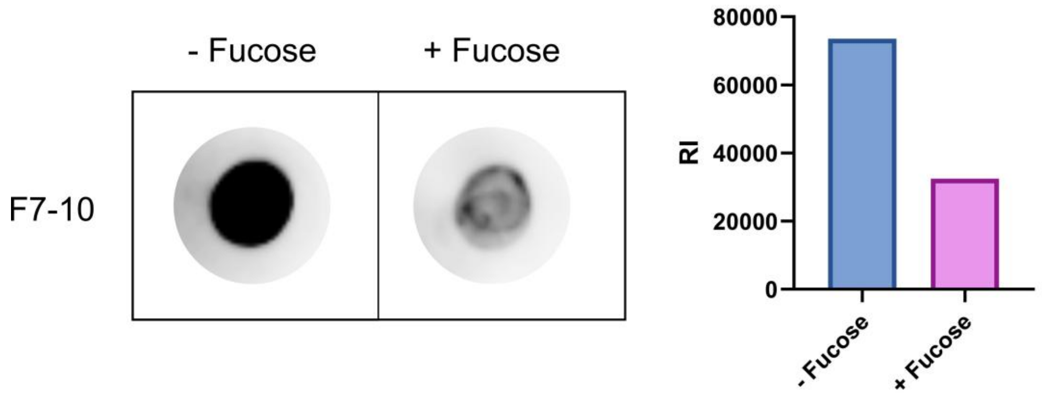


CTxb immunoblot of EVs (F7-10) isolated from *A. suum* by SEC in the presence or absence of L-fucose with densitometric quantification of the immunoblot.

**Supplemental Figure 4**

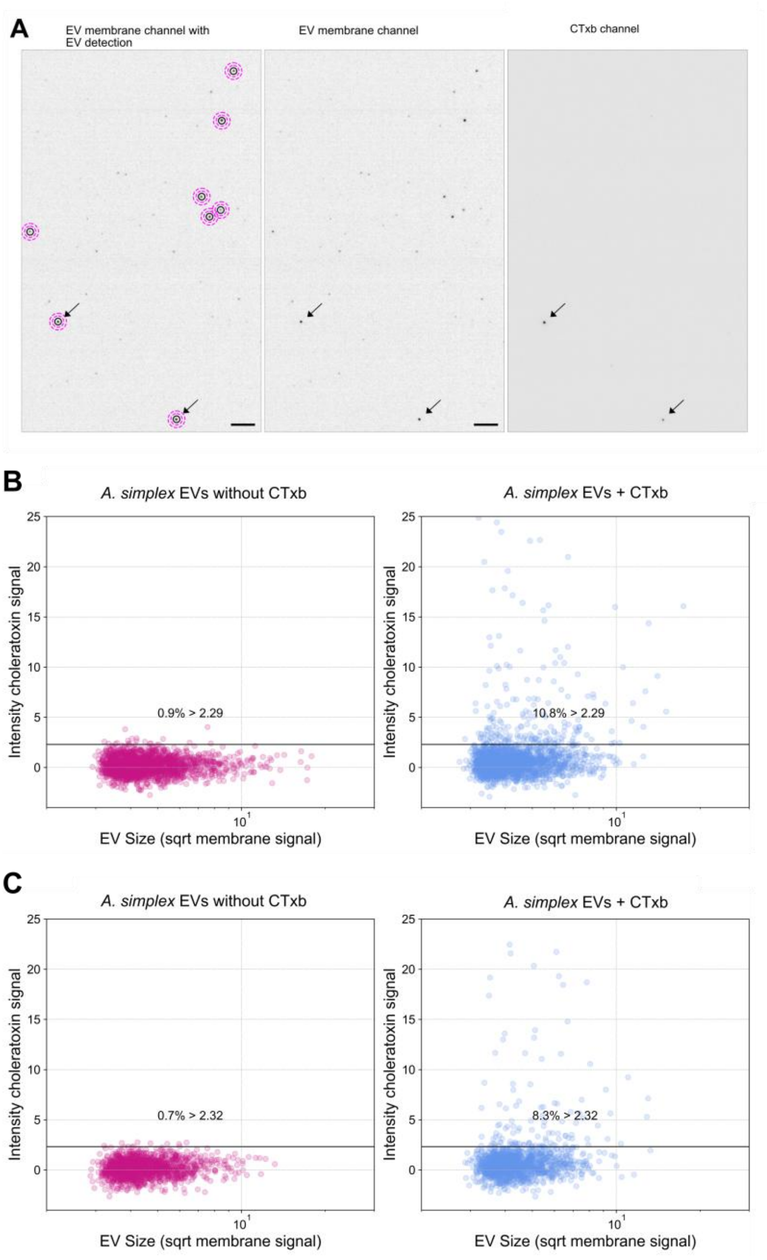


**A:** Representative image showing the co-localization of EVs with CTxb. Left: EVs detected by automated image analysis, the area within the black circles was integrated to obtain the fluorescence intensity of the spots in both channles. The area between the dotted pink circles was substracted for background correction of each spot. Arrows indicate CTxb-positive EVs. Scale bar: 10 µm. **B,C)** The two other individual experiments used in the quantification of CTxb-positive *Anisakis* spp. EVs.

### Supplemental Figure 5

| Protein Description | *Ascaris suum* | *Trichuris muris* | *Brugia malayi* | *S. mansoni* | Min. seq. identity (%) |
|---|---|---|---|---|---|
| Glyceraldehyde 3-phosphate dehydrogenase (GAPDH) | TrEMBL\|U1M303\|Release | TMUE_s0094001000 | XP_001899849.1 | - | 73.24 |
| Small heat shock protein (SHSP) domain-containing protein | UPI000396C4CC | TMUE_s0203001300 | XP_001894373.1 | - | 34.56 |
| SHSP domain-containing protein | UPI000396C4CC | TMUE_s0023008400 | XP_001894373.1 | - | 23.40 |
| Malate dehydrogenase | UPI0003965025 | TMUE_s0005004200 | CRZ22215.1 | - | 62.73 |
| Enolase | TrEMBL\|U1LSA2\|Release | TMUE_s0102000900 | CRZ22135.1 | Q27877.1 | 71.43 |
| Galectin | tr\|F1LAD2\|F1LAD2_ASCSU | TMUE_s0119002100 | XP_001900812.1<br>XP_001896448.1 | - | 40.96 |
| SHSP domain-containing protein | UPI000396C4CC<br>TrEMBL\|U1LZJ4\|Release | TMUE_s0039004700 | XP_001894373.1<br>XP_001894373.1 | - | 23.26 |
| HSP90 | TrEMBL\|U1MCU7\|Release | TMUE_s0014013200 | CRZ24513.1 | - | 75.10 |

EV-proteins with orthologues present in EVs from the *Ascaris suum, Trichuris muris,* and *Brugia malayi*.

### Supplemental Figure 6

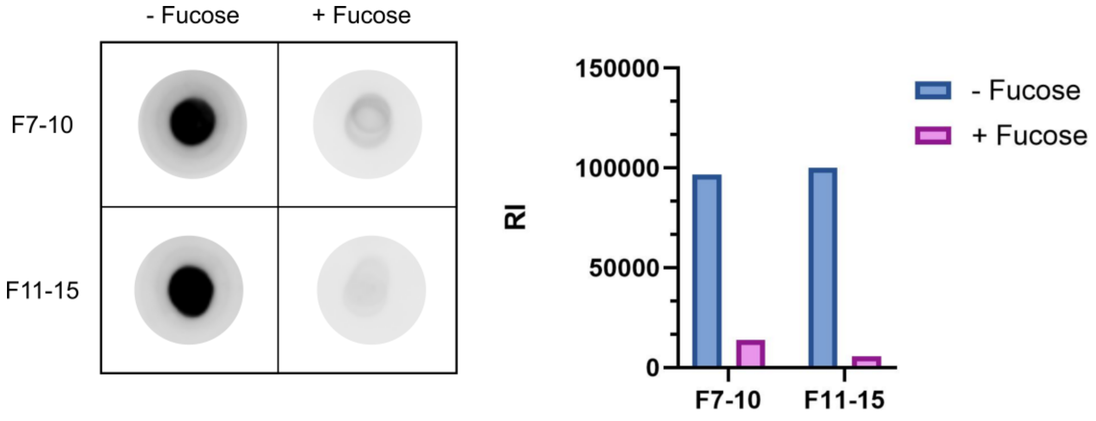


UEA-I lectin immunoblot of EVs (F7-10) and proteins (F11-15) isolated from *A. suum* by SEC in the presence or absence of L-fucose with densitometric quantification of the immunoblot.
